# Database-independent analysis of probable post-translational modifications of soil proteins across ecosystems

**DOI:** 10.1101/2022.02.24.481781

**Authors:** Robert Starke, Stephanie Serena Schäpe, Tim van den Bossche, Tijana Martinovic, Maysa Lima Parente Fernandes, Manuel Delgado-Baquerizo, Felipe Bastida, Nico Jehmlich

## Abstract

The identification rate of measured peptide spectra to proteins barely scratches 1% in best-case scenarios. Hundreds of thousands of valuable spectra are lost as no viable match in the database is found. Here, we apply the delta m/z plot that was previously implemented in MSnbase as tool for quality control to 63 soil samples from three ecosystems with different vegetation (39 forests, 11 grasslands, and 13 shrublands) with the aim to extract probable post-translational modifications (PTM) without the need of a reference database. The validity of the approach was verified with amino acids proposed for their respective 1 Da mass interval and compared to their relative abundance in proteins. We found that the average probable PTM and most known PTMs proposed for the mass intervals are similar across ecosystems. Otherwise, 11 mass intervals changed significantly in relative abundance in the three ecosystems but only for one an annotation could be proposed. Our approach not only highlights the opportunity of the database-independent analysis in soil metaproteomics but paves the way for targeted analysis of the yet unknown PTMs.

---

Among the culture-independent “omic” techniques deployed to gain deeper insights into the structure and function of microbial communities [1], metaproteomics has gained more and more interest from the scientific community as a central element in microbial ecology studies [2], since it deciphers the functional relationships between community members [3]. The workflow involved the extraction of proteins, their tryptic digestion to peptides that are separated with liquid chromatography and measured with mass spectrometry, followed by the identification and annotation of the features with a database of protein coding sequences [4]. In the peptide spectrum matching, the best hit is selected from the database and reported with a matching score that differs from algorithm to algorithm and a false discovery rate (FDR) based on the target decoy of the provided database [5]. The chances for peptide matching increase with the knowledge of what is present in the sample as tailor-made database can be provided [4]. Otherwise, all known protein coding sequences from UniProtKB/SwissProt can be used, however, not only does it contain mostly bacterial sequences, but the ever increasing size leads to an overestimation of FDRs [6] and the loss of valuable protein identification [7]. We expect the loss of much of the valuable information due to incomplete databases, particularly in soils with billions of organisms belonging to thousands of different species per gram [8]. Here, we aimed to use a database-independent approach using the delta m/z plots (the modified *R* script is available in the supplementary information) published as quality control tool [9] and implemented in MSnbase [10] to identify probable PTMs to get a first glimpse of different soil metaproteomic samples without annotation. Delta m/z plots identify the mass differences of peaks in a mass window and could thus be used to identify probable PTMs. For this, we extracted the delta m/z plots using the top 10% most abundant peaks from raw data of 63 soil samples from four continents with different vegetation types (39 forests, 11 grasslands, and 13 shrublands). These samples have been previously used to describe the microbial community from archaea [11], bacteria [12], and fungi (in revision) using reference databases. We estimated the average probable PTM mass between 10 and 250 Da and proposed chemical identities to 1 Da windows. Amino acids were used for verification and known PTMs for targeted analysis. Significant differences of m/z windows between the biomes were estimated by the Kruskal-Wallis test [13]. We hypothesized the presence of probable PTMs specific to each ecosystem and that the average delta m/z is lowest in forests due to the highest microbial activity and its frequent turnover of protein.

Even though we searched in only a small mass window (10-250 Da), we identified thousands of different delta m/z, which is why we condensed the information to 1 Da windows. This removes correctly identified PTMs but it makes the data easier to visualize and interpret. The delta m/z plots showed global differences between the three ecosystems as forests and grasslands comprised a higher contribution of low delta m/z peaks while shrublands showed a higher number of high ones (**Figure 1a**). This trend is also reflected by the average probable PTM, however, without a significant difference (**Figure 1b**), opposite to the protein richness that peaks in forests and grasslands [11,12,14]. Supposedly, shrublands contain less proteins with higher and heavier modifications. Our approach of probable PTMs by delta m/z plots was verified as the relative abundances of the probable amino acids in the delta m/z plot were compared to the relative abundances of amino acids in proteins, showing a high degree of similarity for most of the amino acids (**Figure 1c**). Even though the delta m/z data also contains modifications other than these amino acids in their 1 Da window, only the isomeric amino acids leucine and isoleucine (both 131.175 Da) as well as glutamine (146.146 Da) and lysine (146.189) had relative abundances much below the expected value. While removing correctly identified PTMs by compressing the data to 1 Da windows, this approach worked well for most amino acids, indicating that there might only be one important PTM per 1 Da window an average. Indeed, other modifications with 146 Da difference include the loss of fucose. Interestingly, the relative abundances of the probable PTMs were similar to ∼10% of the relative abundance of amino acids in proteins, which could mean that only 10% of the peaks are actually belonging to proteins that, in turn, could explain the low identification rates in soil metaproteomics. All amino acids besides glutamate showed statistically indifferent relative abundances in the three biomes. Unsurprisingly, known PTMs proposed for the delta m/z windows were similarly abundant in the three biomes with methylation as most abundant followed by dimethylation and oxidation; these modifications are necessary for protein function and should be highly abundant in every soil sample (**Figure 2a**). Only one, glutamate (E), of the abundant significant delta m/z windows could be proposed with a chemical identity while the other ten significant modifications across the three biomes remained without one (**Figure 2b**), which highlighted that yet unidentified PTMs, the dark matter PTMs, could provide a reasonable avenue of exploration when it comes to the differentiation of soil biomes using metaproteomics since known PTMs are likely abundant everywhere.

**Figure 1:**
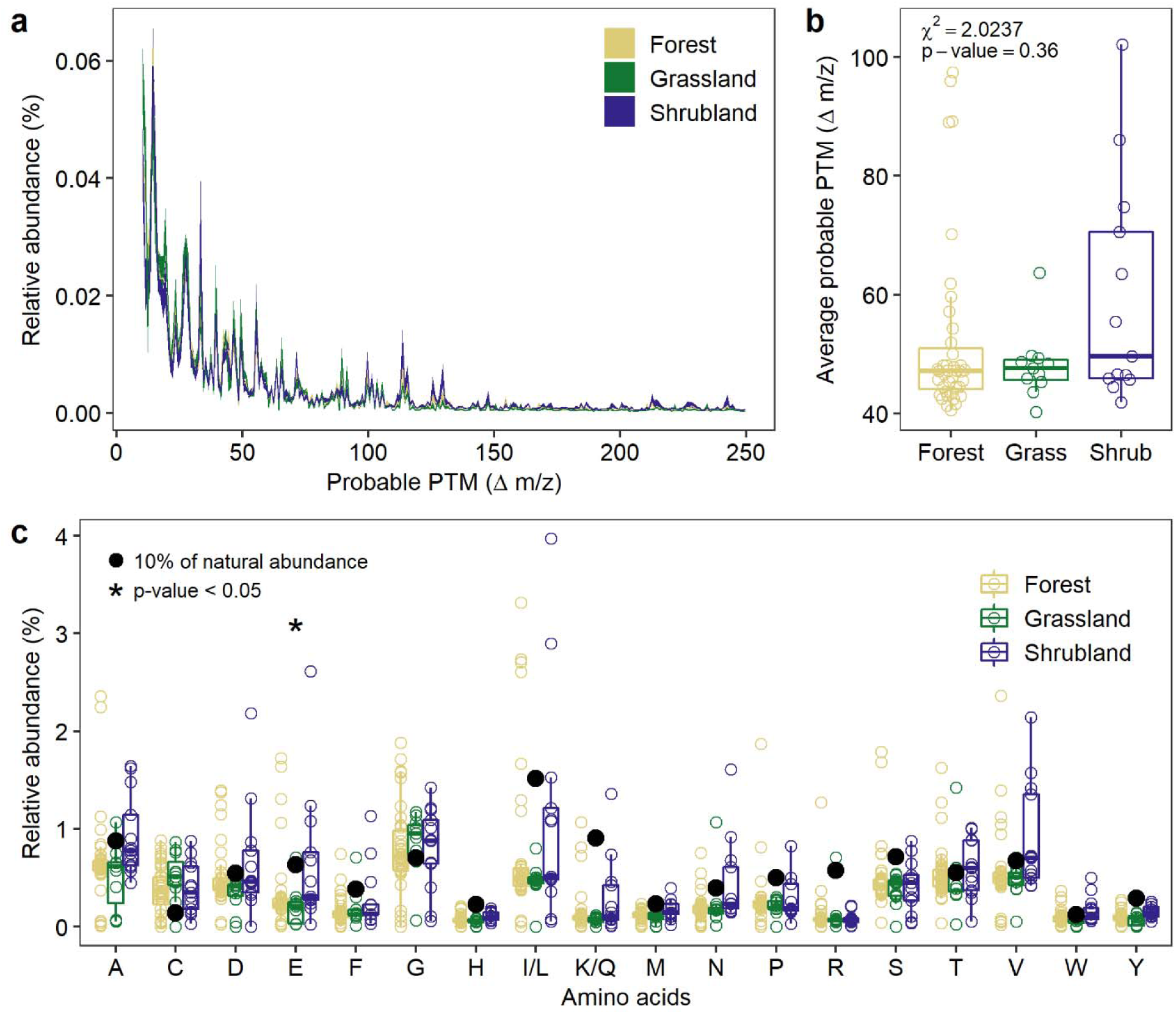
Line plots of probable post-translational modifications with a mass difference between 10 to 250 m/z using the top 10% peaks in every spectrum depending on the biome as line plot with standard error (a). Average probable PTM across the three biomes with the results from the Kruskal-Wallis test for significant differences (b). Box plots of relative abundances of amino acids in the 1 Da window of the delta m/z in each biome (c). Significant differences from the Kruskal-Wallis test and 10% of the average relative abundance of amino acids in proteins are indicated.

**Figure 2:**
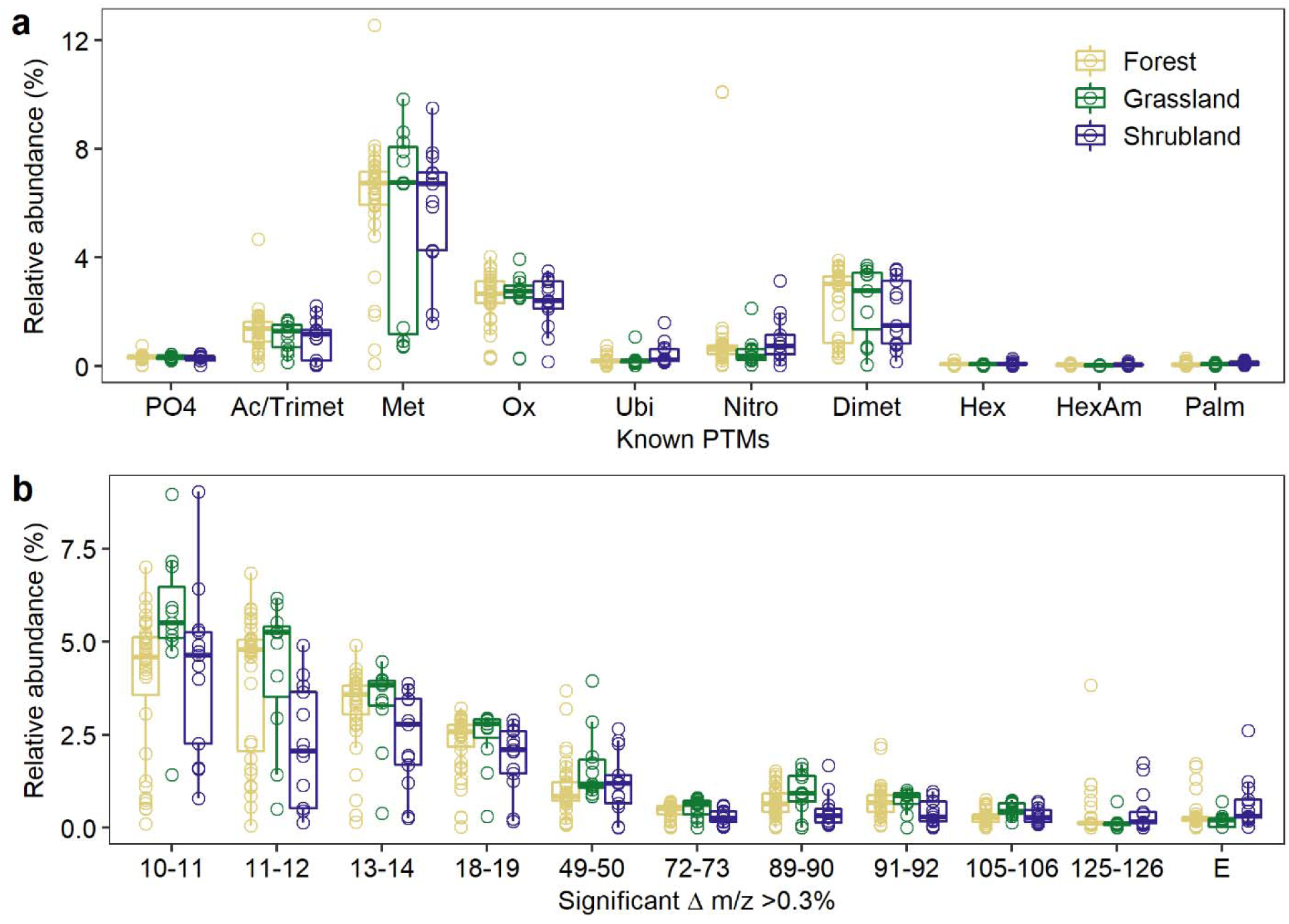
Box plots of relative abundances of known PTMs in the 1 Da window of the delta m/z in each biome (a). No significant differences were found with the Kruskal-Wallis test. PO4 stands for phosphorylation, Ac for acetylation, Trimet for trimethylation, Met for methylation, Ox for oxidation, Ubi for ubiquitinoylation, Nitro for nitrosylation, Dimet for dimethylation, Hex for hexosylation, HexAm for hexosaminylation, and Palm for palmoylation. Box plots of relative abundances of 1 Da windows of the delta m/z with significant differences estimated by the Kruskal-Wallis test in each biome (b). Glutamic acid (E) was proposed for the delta m/z from 129 to 130.

Together, we provided a database-independent way of data analysis in soil metaproteomics that was verified by the relative abundance of amino acids in the data compared to the relative abundance in proteins. Critically, the chemical identities in this study are probable and need to be verified for which protein identification is necessary. The low contribution of amino acids to the abundant peaks in the spectra could explain the low peptide identification rates commonly found in soil metaproteomics and highlight the need of better protein extraction and separation methods as too much of the sample appears to be non-proteinaceous.

## Acknowledgements

Authors thank the Spanish Ministry and FEDER funds for the project AGL2017–85755-R, i-LINK+ 2018 (LINKA20069) from CSIC, and funds from “Fundación Séneca” from Murcia Province (19896/GERM/15). We would like to thank the researchers involved in the CLIMIFUN project for the help with soil sampling. This project has received funding from the European Union’s Horizon 2020 research and innovation program under the Marie Sklodowska-Curie grant agreement No 702057. M.D-B. is supported by a Ramón y Cajal grant from the Spanish Government (agreement no. RYC2018-025483-I). RS thanks Laurent Gatto for modifying the *R* script as well as Alexandra Elbakyan for accessing literature.

## Compliance with ethical standards

The authors declare that they have no conflict of interest.

